# Classification of naturally evoked compound action potentials in peripheral nerve spatiotemporal recordings

**DOI:** 10.1101/469874

**Authors:** Ryan G. L. Koh, Adrian I. Nachman, José Zariffa

**Author notes:** Correspondence and requests for materials should be addressed to José Zariffa.

## Abstract

Peripheral neural signals have the potential to provide the necessary motor, sensory or autonomic information for robust control in many neuroprosthetic and neuromodulation applications. However, developing methods to recover information encoded in these signals is a significant challenge. We introduce the idea of using spatiotemporal signatures extracted from multi-contact nerve cuff electrode recordings to classify naturally evoked compound action potentials (CAP). 9 Long-Evan rats were implanted with a 56-channel nerve cuff on the sciatic nerve. Afferent activity was selectively evoked in the different fascicles of the sciatic nerve (tibial, peroneal, sural) using mechano-sensory stimuli. Spatiotemporal signatures of recorded CAPs were used to train three different classifiers. Performance was measured based on the classification accuracy, F_1_-score, and the ability to reconstruct original firing rates of neural pathways. The mean classification accuracies, for a 3-class problem, for the best performing classifier was 0.686 ± 0.126 and corresponding mean F_1_-score was 0.605 ± 0.212. The mean Pearson correlation coefficients between the original firing rates and estimated firing rates found for the best classifier was 0.728 ± 0.276. The proposed method demonstrates the possibility of classifying individual naturally evoked CAPs in peripheral neural signals recorded from extraneural electrodes, allowing for more precise control signals in neuroprosthetic applications.

## 1. Introduction

The development of neural interfaces for recording peripheral nerve activity, including signal processing algorithms to analyze this activity, is a rapidly growing area of research [1-7]. Peripheral neural signals have the potential to provide the necessary motor, sensory, or autonomic information for robust control in many neuroprosthetic [8-12] and neuromodulation applications [13-18].

However, there are two major challenges in unlocking the potential of these applications.

1. Using an appropriate neural interface to record the neural signals reliably.
2. Developing an algorithm that can correctly recover the information encoded in these signals in order to translate them into command signals.

Of interest here is the task of selective recording, which refers to the ability of a neural interface to discriminate between the activity of different neural pathways. We use the term “neural pathway” to refer to a group of neighbouring nerve fibers with a particular function (e.g. a command signal to a muscle, sensory input from a region of interest, etc.).

Peripheral nerve interfaces to date have had limited success in implementing selective recording strategies that are suitable for long-term use in humans. Intraneural electrodes that penetrate the nerve, such as longitudinal or transverse intrafascicular electrodes or microelectrode arrays, are able to selectively record neural activity and are increasingly being translated to human studies [10, 19, 20] but still struggle to demonstrate viability for stable chronic recordings. Extraneural electrodes, such as nerve cuffs or flat interface nerve electrodes (FINEs), have been shown to be stable for chronic implantation in humans for recording [21-22] and stimulation [23-25] applications. Thus nerve cuff electrodes are an appealing choice of neural interface, but it is a challenge to achieve sufficient recording selectivity with these devices.

Several different algorithms have been developed to try to improve the recording selectivity of extraneural electrodes using multi-contact configurations. These techniques revolve around two general approaches: identifying conduction velocity (temporal information), or identifying the location of the signal source within the nerve (spatial information). In the first category, velocity-selective recording (VSR) methods [26-28] have taken advantage of action potential propagation via delay-and-add operators that emphasize compound action potentials (CAP) with specific velocities and make it possible to infer the fiber type. However, this approach is unable to distinguish signals that have similar conduction velocities. In the second category, source localization approaches [29-31] have been used with some success, and some algorithms following a beamforming approach [7, 32-34] have shown the ability to distinguish signal sources adequately. However, the performance of these methods can degrade rapidly with poor signal-to-noise ratios (SNR), often calling for rectify-bin-integration (RBI) or other windowing techniques that lower the temporal resolution. Building on these ideas, our previous work [5] has shown that recording selectivity can be improved by integrating spatial and temporal information together into spatiotemporal templates, thus more fully characterizing each neural pathway of interest.

Most of the studies mentioned above involved CAPs that were evoked electrically using stimulation electrodes [26-28, 30-33], leading to much higher amplitude and less physiologically realistic CAPs compared to naturally evoked CAPs (e.g. produced by proprioceptive or mechano-sensory afferent activity). For studies involving naturally evoked afferent activity [7, 34], classification was applied to windowed signals (e.g. rectify-bin-integrated), not to individual CAPs. Only [28, 35] have attempted clustering of naturally evoked CAPs by their velocities. However, this technique cannot deal with neural activities which have similar conduction velocities. [36] demonstrated clustering of natural CAPs and associated cluster firing rates with different physiological events. To date, no study has successfully classified *individual, naturally evoked CAPs* in order to discriminate the activity of neural pathways of interest.

We refer to the task of identifying the signal source of an individual detected CAP as CAP-based classification. Note that naturally-evoked CAPs may be referred to as “spikes” in the nerve cuff literature, but we use the term CAP here to avoid confusion with single-fiber activity, which is expected to be well below the noise floor in a nerve cuff recording [37].

CAP-based classification is made difficult by the low SNR, but if successful would have important advantages. First, the firing rates of individual pathways could be reconstructed with a finer time resolution. In state of the art techniques, RBI windows used can range anywhere from 33ms [34] to 100ms [7, 32, 33], whereas classifying individual CAPs would result in a time resolution of under 5 ms. Second, it could improve selective recording performance in situations where multiple neural pathways are active over a given period of time. This second advantage comes about because averaging data over time intervals increases the likelihood that a source localization or beamforming approach would need to deal with multiple simultaneous signal sources. In contrast, the probability that individual CAPs will overlap in time is much lower, resulting in an easier source identification problem. In our previous simulation study [5], we used spatiotemporal templates to interpret peripheral nerve activity recorded with multi-contact nerve cuff electrodes, using a CAP-based classification framework. The rationale for this approach was that the templates could more fully characterize the neural signals, as well as leverage the redundancy afforded by multiple rings of contacts to help deal with the low SNR.

Here, *in vivo* CAP-based classification is investigated. The simulation results from our previous study [5] are validated *in vivo*, and alternative classification schemes are compared in order to maximize the CAP-based classification performance that can be achieved using spatiotemporal signatures extracted from multi-contact nerve cuff recordings. The main contribution of this work is to demonstrate for the first time the classification of *individual, naturally evoked CAPs* in extraneural peripheral nerve recordings, using spatiotemporal signatures.

## 2. Methods

### 2.1 Experimental Protocol

Acute experiments were performed on 9 Long-Evans rats (retired breeders) under isoflurane anesthesia. The lower back and legs were shaved and treated with povidone-iodine. Once an adequate level of anesthesia was reached (indicated by loss of response to a sharp toe pinch), the animals were positioned prone on the operating table.

An oblique incision was made on the posterior and dorsal aspect of the hip as shown in Figure 1a. The incision provided direct access to the sciatic nerve through the natural splitting of the fibers of the gluteus maximus while clearing the skin and deep fascia. The sciatic nerve was exposed as far proximally as possible to allow for the application of the nerve cuff electrode. Figure 1b shows the implanted nerve cuff electrode on one of the rats.

**Figure 1.**
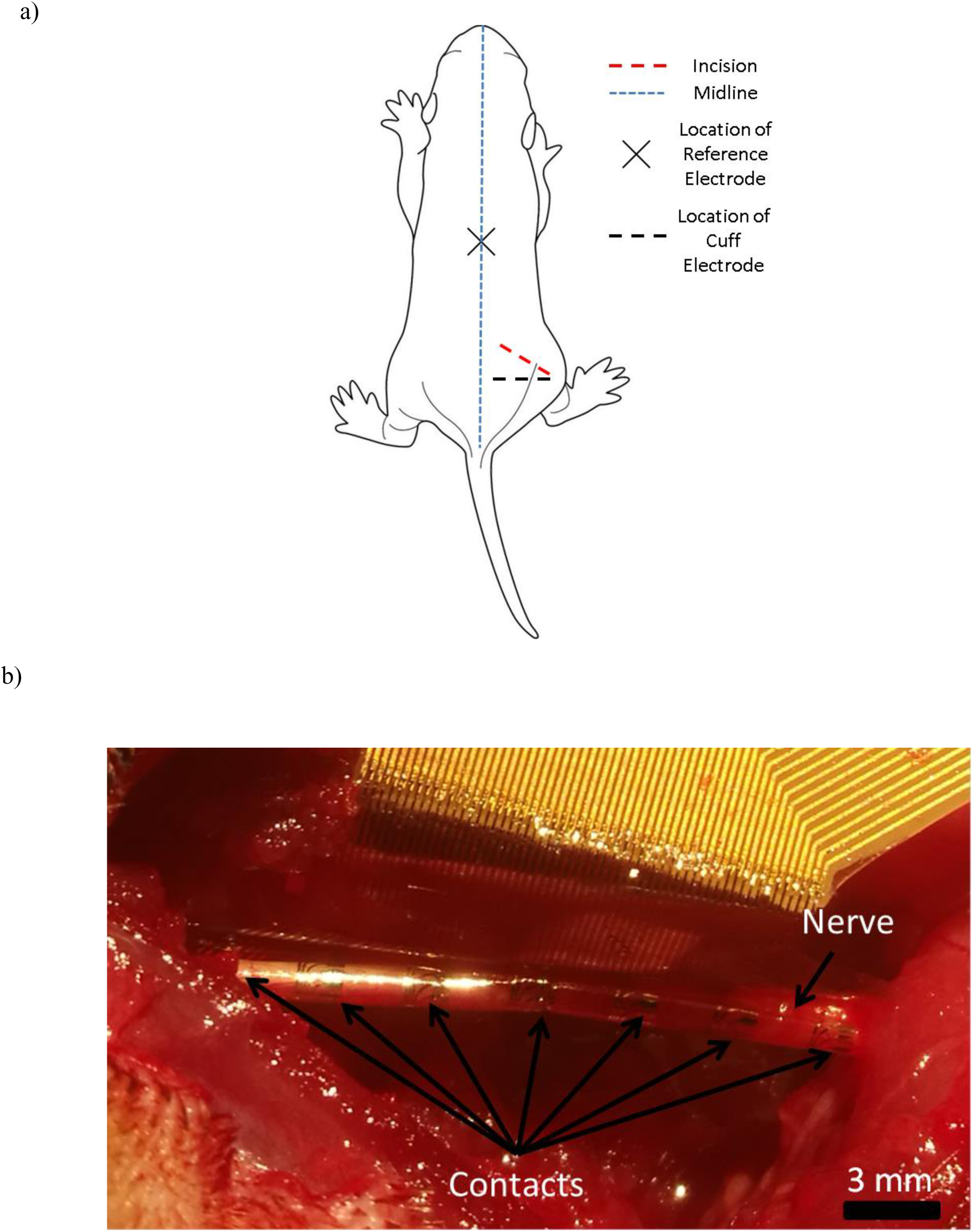
(a) The oblique incision position on the rat, and the location of the needle electrode used as a reference and the location of the nerve cuff electrode. (b) The implanted nerve cuff electrode on the sciatic nerve.

Neural activity was recorded using a 56-multi-contact spiral polyimide nerve cuff electrode (CorTec GmbH, Freiburg, Germany). The nerve cuff consisted of 7 rings of 8 contacts distributed over the length of the cuff electrode. The cuff had a length of 23 mm and a diameter of 1 mm. A needle electrode was placed in the back of the animal to serve as a reference. Data was acquired using a neural data acquisition board (RHD2000, Intan Technologies, USA) with a sampling frequency of 30 kHz and the recordings were bandpass filtered on the acquisition board between 256 Hz and 7.5 kHz.

Afferent activity was selectively evoked in the different fascicles of the sciatic nerve (tibial, peroneal, sural) using mechanical stimuli. The foot was held by the claws using a clip with plastic tips and the ankle was manually dorsiflexed and plantarflexed by approximately 60° to evoke proprioceptive activity in the tibial and peroneal branches, respectively [38]. The ankle was moved to be in sync with a metronome set to a tempo of 70 beats per minute. At the 1^st^ beat, the ankle was moved to the maximum dorsiflexion/plantarflexion point (∼60º), held in place on the 2^nd^ beat and moved back to the initial position on the 3^rd^ beat and this process was repeated until 100 trials were recorded. A cutaneous stimulus to the heel was used to elicit activity in the sural branch [39] using a Von Frey monofilament (300g). Following the metronome, the 1^st^ beat involved pricking of the heel and held in place until the 2^nd^ beat, when the Von Frey was removed from the heel and this process was repeated until 100 trials were reached. A combined stimulus, involving the alternation between dorsiflexion and plantarflexion, was also elicited 100 trials were performed for each stimulus. An example of the neural activity is shown in Figure 2.

**Figure 2.**
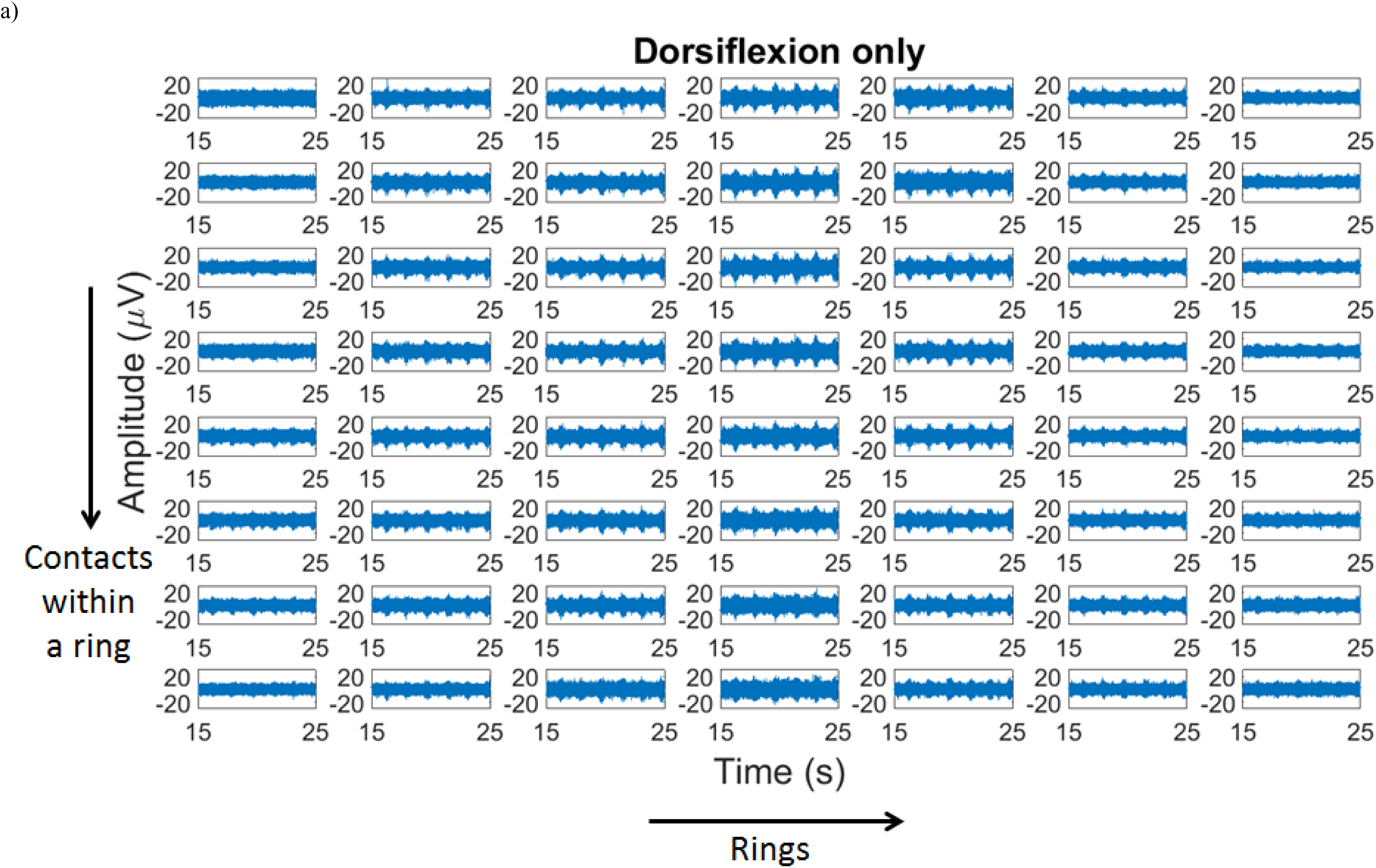
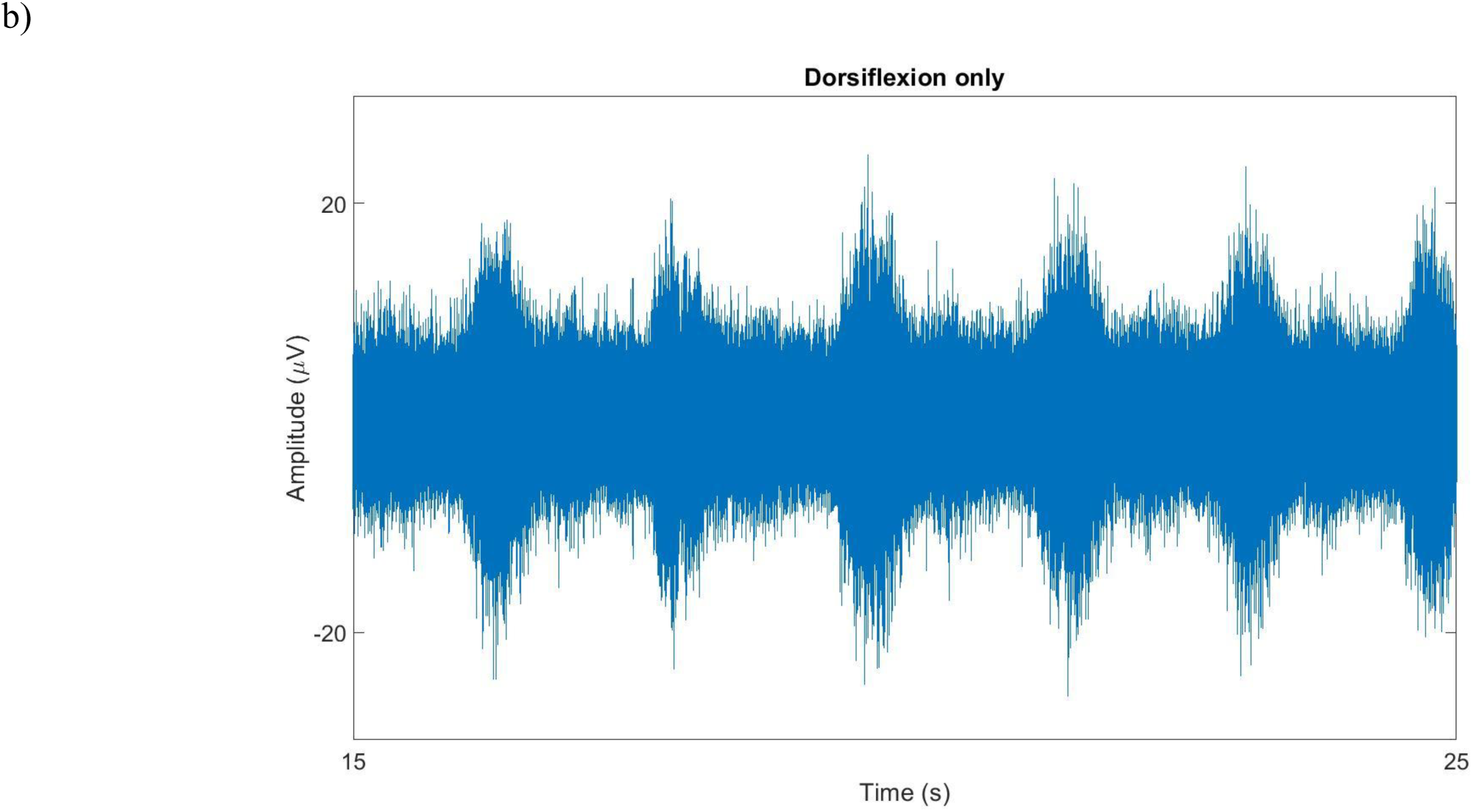
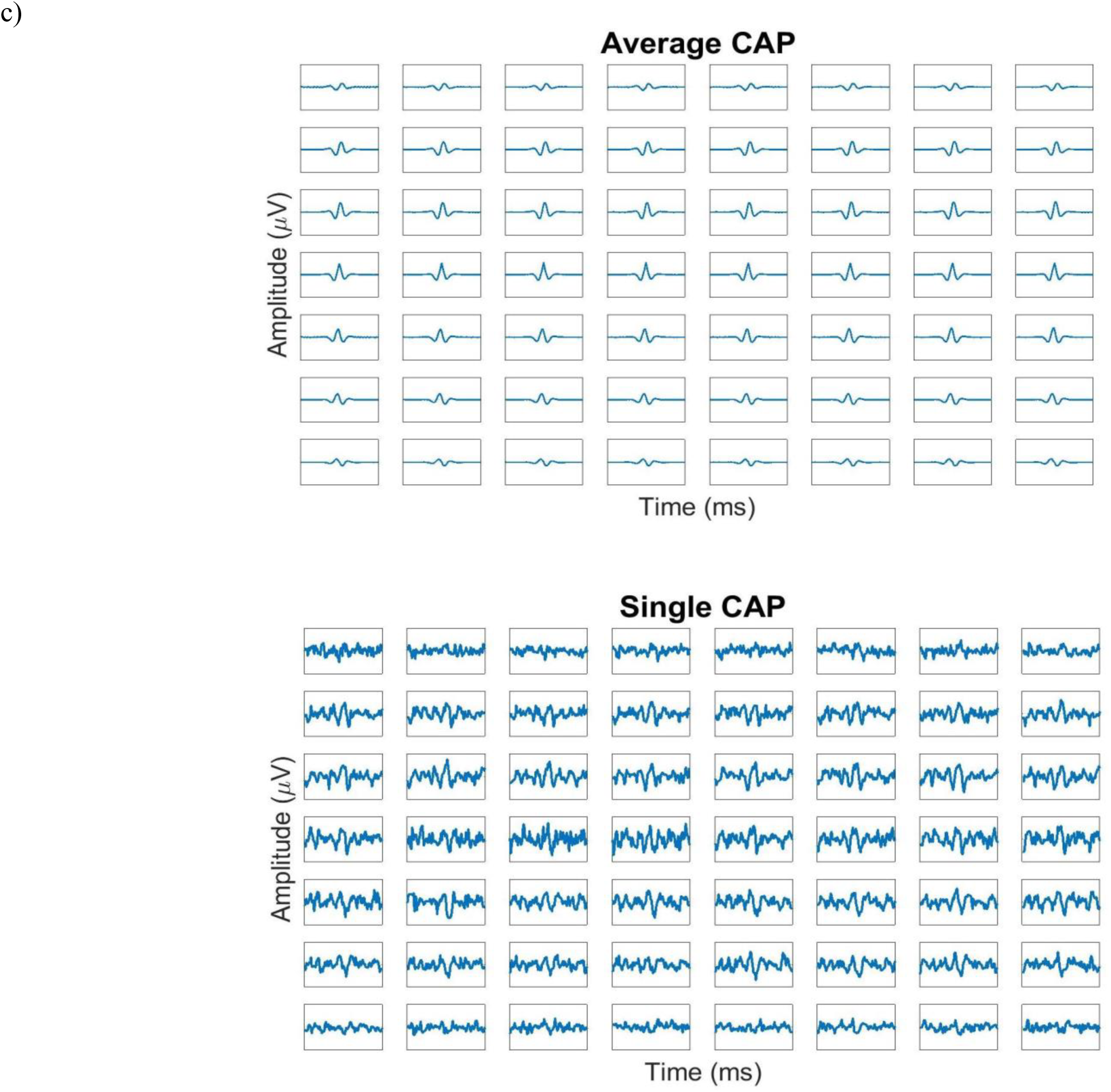
(a) Example of the neural recording from dorsiflexion only at all contacts in Rat 6. The grid corresponds to 7 rings, each consisting of 8 contacts (b) Plot of a middle ring contact’s neural recording. (c) Recorded signal at all contacts for the dorsiflexion only stimulus. Top - CAP recorded in all contacts, averaged over 5788 trials, Bottom - Single detected CAP in all contacts. All boxes are from 0 – 3.33 ms and amplitudes from -20 to 20 μV.

The applications of the stimuli were video recorded, synchronized with the neural recordings through a LED visible in the video, and later manually annotated to determine the times at which each type of stimulus was being applied. All CAPs detected during the application of given stimulus were inferred to have originated from the corresponding neural pathway.

The experimental procedures were approved by the Animal Care Committee of the University of Toronto.

### 2.2 Spatiotemporal Signatures

Spatiotemporal signatures for each activity (dorsiflexion, plantarflexion and pricking of the heel) were constructed using the detected CAPs for each stimulus using *M* contacts at *T* consecutive time samples. This created an *MxT* matrix which was then reshaped into a column vector constituting the spatiotemporal signature for that CAP. By associating each CAP in a new recording with one of these known signatures, the activities of multiple neural pathways can be discriminated. The spatiotemporal signatures were averaged to form templates (Fig. 3) that were then used to create tailored matched filters (MF). The spatiotemporal signatures were further used as the inputs to machine learning-based classifiers. These different classification methods are described in section 2.4. The spatiotemporal templates from each stimulus type were used to calculate conduction velocities to verify the fiber type.

**Figure 3.**
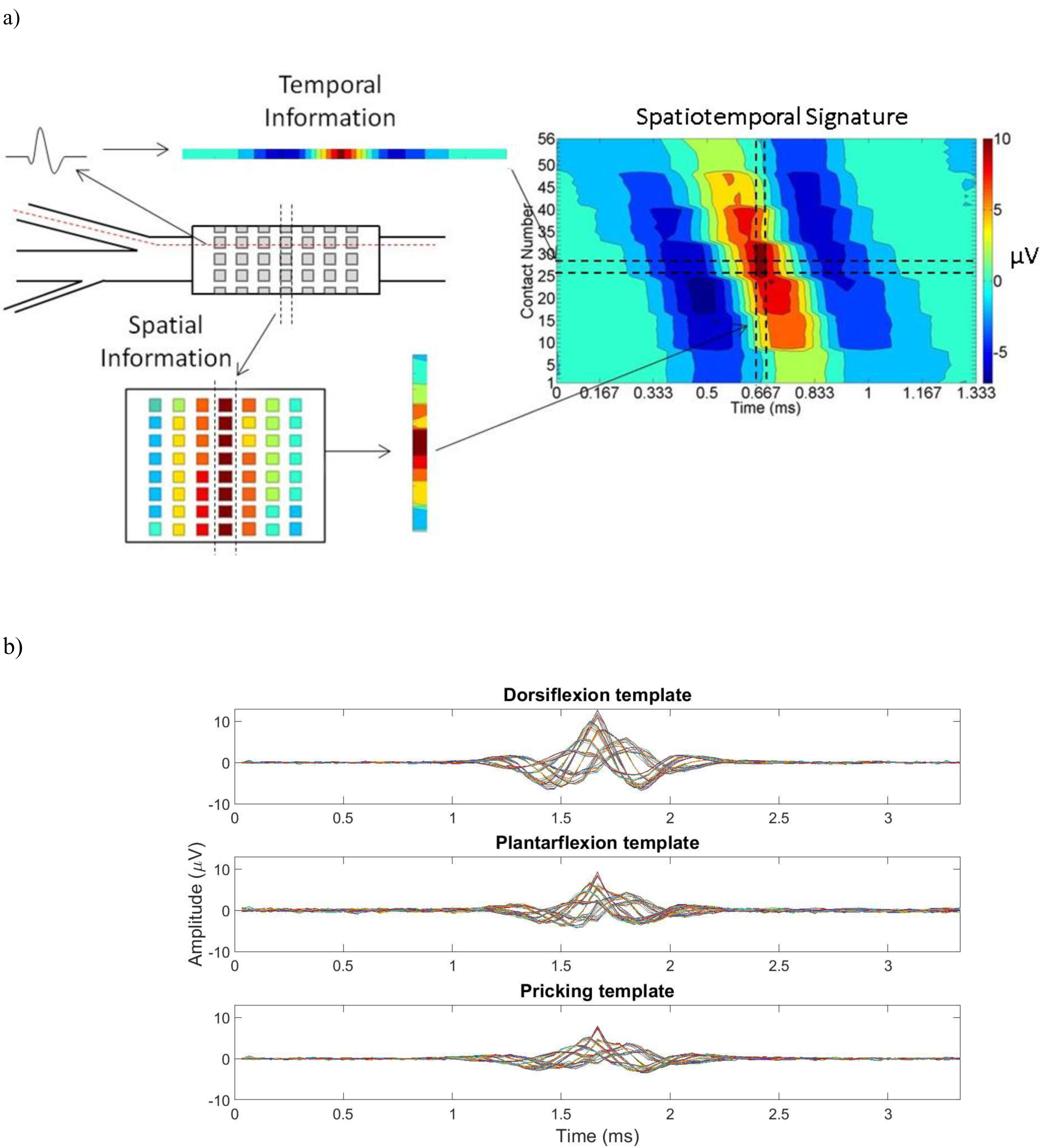
(a) The incorporation of the spatial and temporal information from a given pathway in the peripheral nerve to create a spatiotemporal signature of a specific stimulus. This signature on the right-hand side was created using the average of CAPs recorded during dorsiflexion in one rat. The diagram on the left-hand side illustrates how the signature includes information about the signal over time at individual contacts (top left), as well as the variations in signal over multiple contacts at the same instant (bottom left). The contacts in the bottom left diagram are organized in the same manner as in Figure 2a (b) Example of the average spatiotemporal template of each stimulus type from Rat 6. These templates were created by taking the average of all detected CAPs’ spatiotemporal signatures for a given activity. Each of the 56 signals in a template corresponds to one of the contacts in the cuff. All signals shown in this figure have been tripole referenced with respect to the 2 outer rings.

### 2.3 CAP Detection

The delay-and-add operation used in the velocity selective recording (VSR) method [26-28] was applied to the raw *in vivo* signal after tripole referencing (using the averages of all the contacts in the two outermost rings of the cuff as the reference). This output X was then thresholded using the median absolute deviation estimate approach (Eq. 1) [40] to find CAPs. CAPs with peak values above 15μV were discarded, and the rest of the detected CAPs were then aligned to the peaks of the waveforms in the center ring.

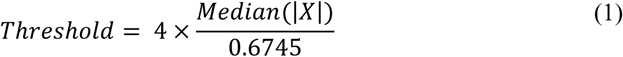

Once the CAPs were detected, 49 time samples before and 50 time samples after the peak were used to create the spatiotemporal signatures.

### 2.4 Classification Algorithms

Four classification techniques were used and compared in this study.

The first method was the one described in our previous simulation work [5]. Briefly, the spatiotemporal templates described in Section 2.2 were used to create a MF for each stimulus. Each template was made from the average of detected CAPs for a given stimuli in the training set, and then applied to the test set. The CAP classification was based on the maximum filter output of the MFs at the detected CAPs after dividing by their corresponding normalization factor **u**_i_.

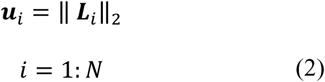

Where **L**_i_ is the spatiotemporal template of the i^th^ stimuli.

The next two methods used in this paper were machine learning approaches, which may be able to uncover additional identifying patterns in the spatiotemporal signatures.

The second method incorporated the spatiotemporal signatures as input data in a random forest classifiers. This classifier was implemented through TreeBagger in Matlab (2012a). A 200 tree random forest was used while the other parameters were the default parameters set in Matlab’s TreeBagger. The inputs used were MT x 1 vectors corresponding to the reshaped spatiotemporal signature for a given CAP.

The third method also used a machine learning approach, this time with a neural net implemented through the patternnet function in Matlab (2012a). All detected CAPs were reshaped into column vectors of MT x 1 and were used to train the neural net. The architecture of the neural net had 4 hidden layers (2000 x 500 x 100 x 20 nodes) with a softmax output layer to produce a probabilistic output. The maximum number of training epochs was 2000 with a learning parameter of 0.2. An early stopping criteria was that the error in the validation set continuously increased more than a maximum of 8 consecutive times.

A fourth method used windowed data rather than individual CAPs. The purpose of this analysis was to obtain an upper bound against which to compare the CAP-based classification performance. This last method was a random forest classifier which was applied to signals preprocessed using the RBI method. These signals were rectified and then integrated over intervals of 1, 10, 33, or 100 ms, as used in previous studies [32-34, 41]. The RBI technique improves the SNR, but loses the advantage of CAP-based classification. In our analysis, peak points in the RBI series were detected by setting a manual threshold for each rat, and the windows to test were extracted around those peak points. From each window, the 56 peak RBI values (one per contact) were used as inputs to the random forest classifier. In this case, classification accuracy was based on the labels of these peak points rather than of individual CAPs. From this point forward this method will be abbreviated as the RF-RBI approach.

### 2.5 Data Augmentation

Data augmentation was performed to enhance the training of the machine learning classifiers, because the number of CAPs detected in several rats created a class imbalance. To address this problem, new samples were created by generating noisy versions of an averaged CAP. First, the average CAP for each neural pathway of interest in each rat was obtained by averaging all CAPs detected while that pathway was being stimulated. Next, the noise at each contact was characterized by subtracting the average CAP from individual CAPs and fitting a Gaussian distribution to the resulting noise-only data. These Gaussian distributions were then used to create new noisy samples by adding noise sampled from the distribution to the average CAP template.

### 2.6 Training and Evaluation

CAPs detected using the median absolute approach (described in Section 2.3) from each activity (the 100 trials each from dorsiflexion only, plantarflexion only, and pricking only) were split into training and test sets using a 3-fold cross validation approach for each rat. Unless explicitly stated, all training sets for the machine learning classifiers described below incorporated data augmentation. This consisted of adding corresponding augmented CAPs for each neural pathway until there were 10000 training examples (combination of detected and augmented CAPs) of each class. Testing sets for all results only consisted of detected CAPs (no augmented CAPs) for each neural pathway.

For the MF approach, these training sets were used to obtain the average spatiotemporal templates, which were then used to create the MFs for classification. For the machine learning approaches, the spatiotemporal signatures of each CAP in the training set were used to train the 2 classifiers. In the RF-RBI approach, a 4-fold cross validation approach was used, due to the reduced number of training examples after applying the RBI technique. Spatiotemporal signatures for both the MF approach and the machine learning classifiers were created from the 56 channels x 100 time samples (3.33ms long).

Classification accuracy was quantified on an individual CAP basis, except for in the RF-RBI results that were quantified based on the peak points. The classification accuracy and F_1_-score of each activity was calculated. This F_1-_score metric indicates a classifier’s predictive power better than classification accuracy when the classes are imbalanced in the test set. In the multi-class problem, F_1_-score was calculated based on the precision and recall of each class, and the reported F_1_-score is the average of the F_1_-scores for all classes.

Lastly, we sought to quantify the effect of the SNR on the ability to discriminate the different stimuli. In order to answer this question, multiple CAPs from each class (in groups of 2, 3, or 4) were averaged, in order to improve the SNR through trial averaging. The classification accuracy was then evaluated as a function of the CAP SNR, using the methods described above. The training sets used for this portion of the analysis did not incorporate data augmentation, in order to quantify solely the effects of SNR on the classification accuracy. Eight of the nine rats were used, based on the criterion that after averaging groups of 4 CAPs, at least 100 or more CAPs should remain in each class.

### 2.7 Reconstruction of firing rates

The third metric which we investigated was the ability to reconstruct the firing rate of each neural pathway. In this case the training set consisted of all CAPs (all peak points for RF-RBI) in the dorsiflexion only trials and plantarflexion only trials in combination with augmented CAPs for each activity. The classifiers were then evaluated on a test set of data from alternating dorsiflexion and plantarflexion. Detected CAPs in the test set were classified and the firing rates of each pathway (dorsiflexion or plantarflexion) were calculated by convolving the estimated CAP trains with a Gaussian kernel with a standard deviation of 150ms [42]. The true firing rates were determined by placing all CAPs detected during the known activation of a given pathway (based on the synchronized video recordings) in a CAP train, and convolving with the same Gaussian kernel. Pearson correlation coefficients were used to describe the relationship between the true and estimated firing rates. Rat 3’s data was removed from this analysis due to the degrading plantarflexion signal throughout the course of the experiment.

Furthermore, to minimize the number of spurious CAPs caused by noise, a threshold on the outputted probabilities of the machine learning classifiers was used to classify a detected CAP as spurious (i.e., if a CAP was not strongly associated with any of the classes it was considered to be noise). This optimal threshold was chosen empirically for each machine learning classifier. All metrics for this portion of the analysis are reported based only on the CAPs that remained after the removal of the spurious CAPs.

### 2.8 Data Availability

The dataset collected and analysed during this study are available from the corresponding authors on reasonable request.

## 3. Results

### 3.1 Number of Detected Points of Interest and Velocity

The mean and standard deviation of detected CAPs for each activity from all rats were 3,898 ± 2,175, 3,454 ± 2,433, and 3,991 ± 2,145, for dorsiflexion, plantarflexion and pricking respectively. The mean and standard deviation of the number of peak points detected in the RF-RBI approach for each activity were 101 ± 2, 100 ± 1, and 99 ± 2, for dorsiflexion, plantarflexion and pricking respectively. The average conduction velocity determined from the templates of the spatiotemporal templates across all rats for dorsiflexion, plantarflexion and pricking were 70.06 ± 12.26,71.93 ± 16.96, 56.79 ± 10.34 m/s respectively, which is consistent with the conduction velocities of the expected large myelinated fibers.

### 3.2 Classification Accuracy

Figure 4a shows the mean F_1_-scores of the MF approach, random forest, neural net, and the RF-RBI approach for all window sizes in a 3-class discrimination problem (dorsiflexion, plantarflexion and pricking) from all rats. Similarly Figure 4b shows the mean F_1_-scores in the 2-class problem (dorsiflexion and plantarflexion). Figure 4c shows the corresponding confusion matrices for the 3-class problem and Figure 4d shows the confusion matrices for the 2-class problem.

**Figure 4.**
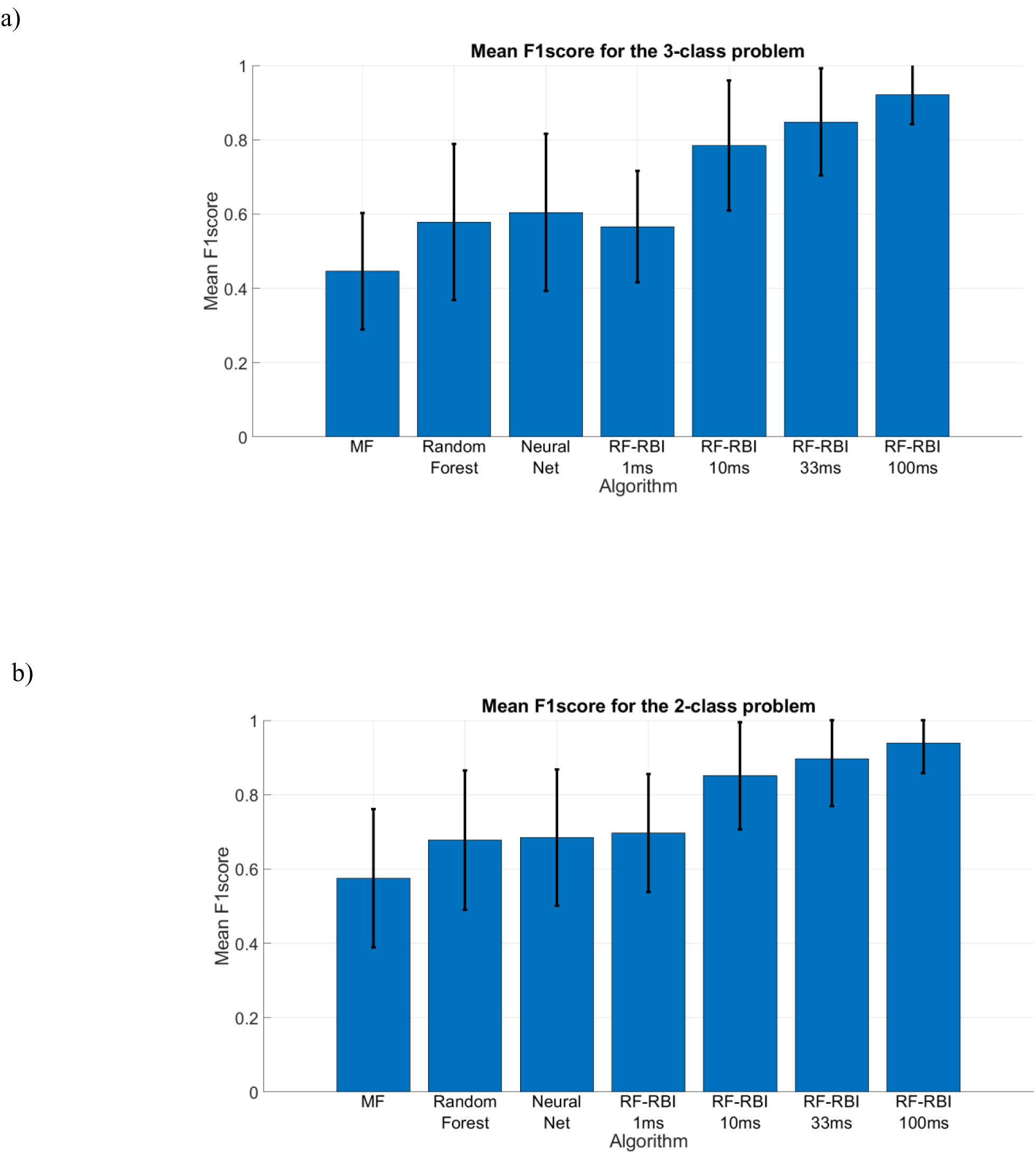
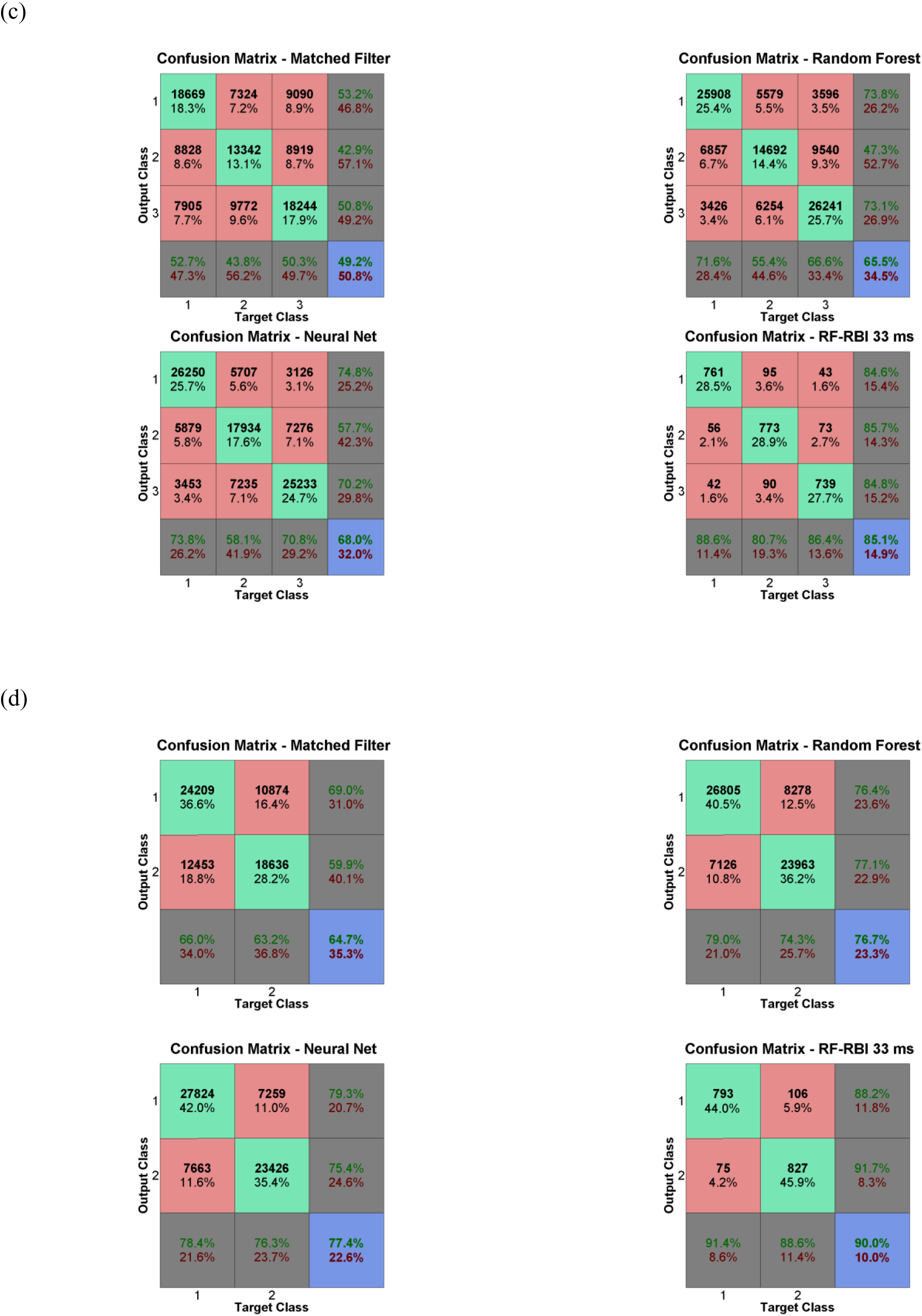
(a) Mean F_1_-score of each algorithm for the 3-class problem (dorsiflexion, plantarflexion, and pricking) (b) Mean F_1_-score of each algorithms for the 2-class problem (dorsiflexion and plantarflexion). (c) Confusion matrices for the different algorithms for the 3-class problem, constructed using combined test sets from all rats. (d) Confusion matrices for the different algorithms for the 2-class problem. Note that the classification accuracies shown in (c) and (d) are slightly different than mean classification accuracies reported in Section 3.2 because they are computed using a combined test set rather than by averaging the classification accuracies for each rat. Class 1 – Dorsiflexion, Class 2 – Plantarflexion, Class 3 – Pricking of the heel.

The mean classification accuracies were 0.510 ± 0.108, 0.658 ± 0.115, 0.686 ± 0.126 and 0.851 ± 0.139 and corresponding mean F_1_-scores were 0.446 ± 0.157, 0.578 ± 0.210, 0.605 ± 0.212 and 0.848 ± 0.144 for the MF, random forest, neural net and RF-RBI with a 33ms window approach respectively for the 3 class-problem. For the 2-class problem the mean classification accuracies were 0.655 ± 0.120, 0.751 ± 0.108, 0.761 ± 0.119 and 0.899 ± 0.123 and corresponding mean F_1_-scores were 0.575 ± 0.186, 0.677 ± 0.188, 0.684 ± 0.183 and 0.897 ± 0.127 the MF, random forest, neural net and RF-RBI with a 33ms window approach respectively.

Additionally, for the alternating dorsiflexion and plantarflexion data, the mean F_1_-scores were 0.619 ± 0.170, 0.614 ± 0.169, and 0.853 ± 0.132 for the random forest, neural net, and the RF-RBI technique with a 33ms window respectively. Figure 5 shows the effects of the SNR on the mean F_1_-score, as a function of the relative improvement from the SNR seen *in vivo* for the classifiers. With a factor of 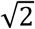 improvement over base SNR levels, the F_1_-scores improve in a 3-class problem by approximately 11%. The general trend in this analysis shows that with increased SNR our ability to discriminate the activity of different neural pathways from multi-contact nerve cuff recordings increases quite substantially.

**Figure 5.**
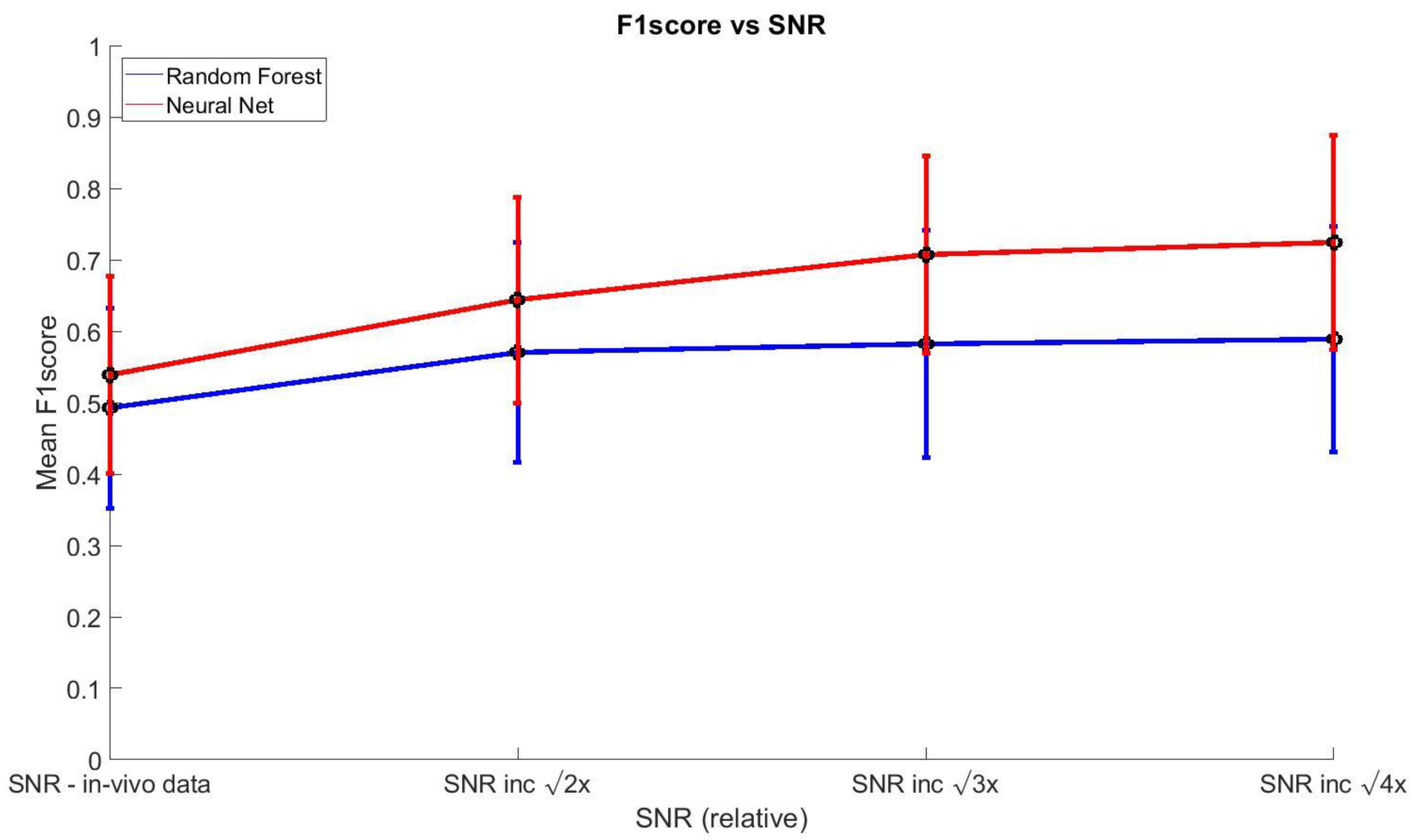
Mean F_1_-score vs SNR improvements relative to SNR observed *in vivo.*

### 3.3 Reconstruction of firing rates

The mean Pearson correlation coefficients were 0.736 ± 0.248, 0.728 ± 0.276,and 0.794 ± 0.158 for the random forest, neural net, and the RF-RBI technique with a 33ms window respectively, for the alternating dorsiflexion and plantarflexion data. The mean Pearson correlation coefficients increased to 0.758 ± 0.218, and to 0.741 ± 0.271, for the random forest and neural net respectively when an optimal threshold was used to identify and remove spurious CAPs. An example of the difference in tracked firing rate with and without removing spurious CAPs can be seen in Figure 6 for Rat 5.

**Figure 6.**
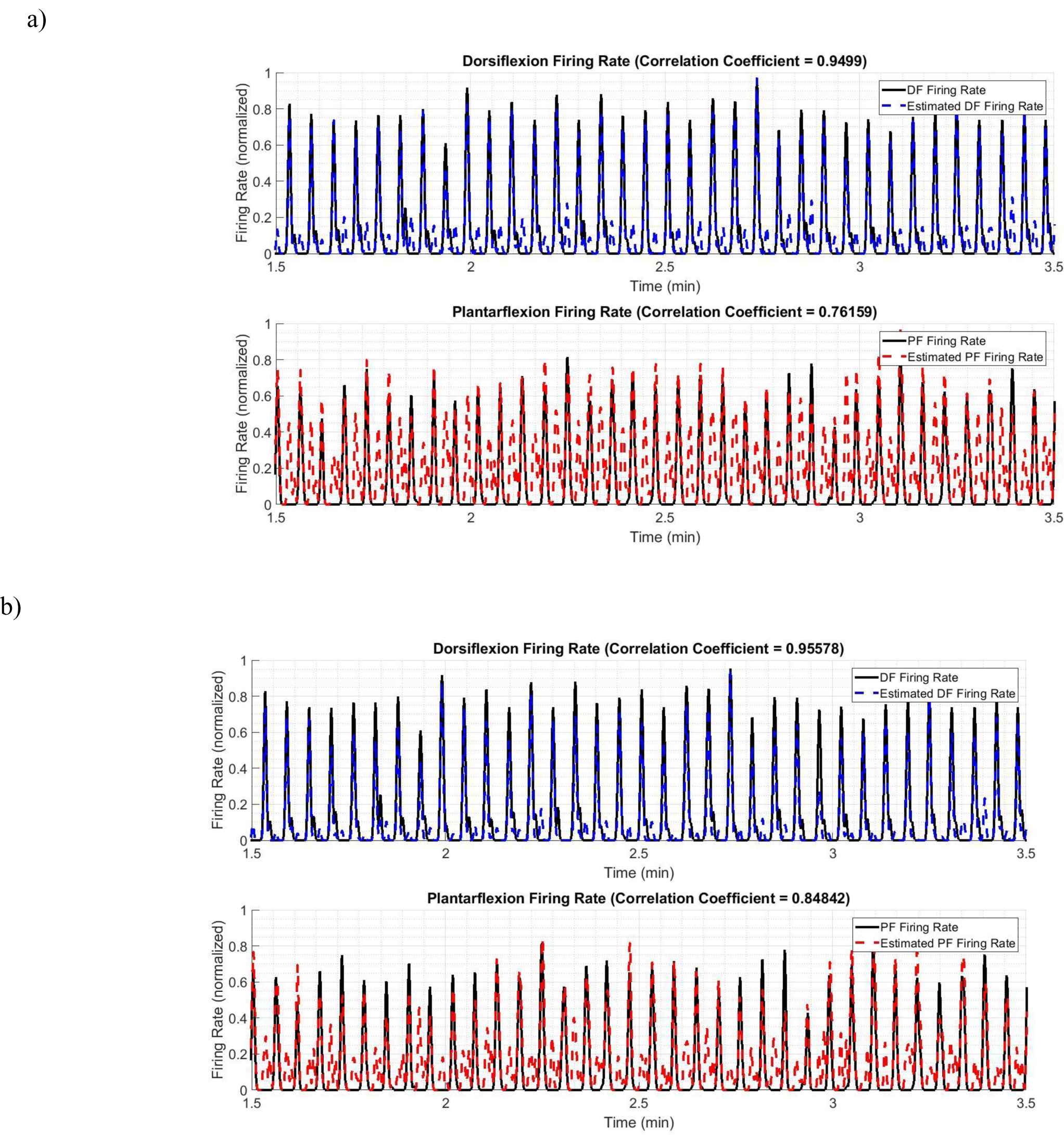
(a) Estimated firing rates using the neural net classifier on CAPs detected by the automatic thresholding method [26]. Mean F_1_-score = 0.686 (b) Estimated firing rates using the neural net classifier on CAPs detected by the automatic thresholding method followed by removal of spurious CAPs using an optimal threshold. Mean F_1_-score = 0.7829. Both plots represent the reconstructed firing rate of Rat 5.

Figure 7 shows the relationship between the ankle angle, tripolar nerve cuff recordings and the reconstructed firing rates from Rat 6.

**Figure 7.**
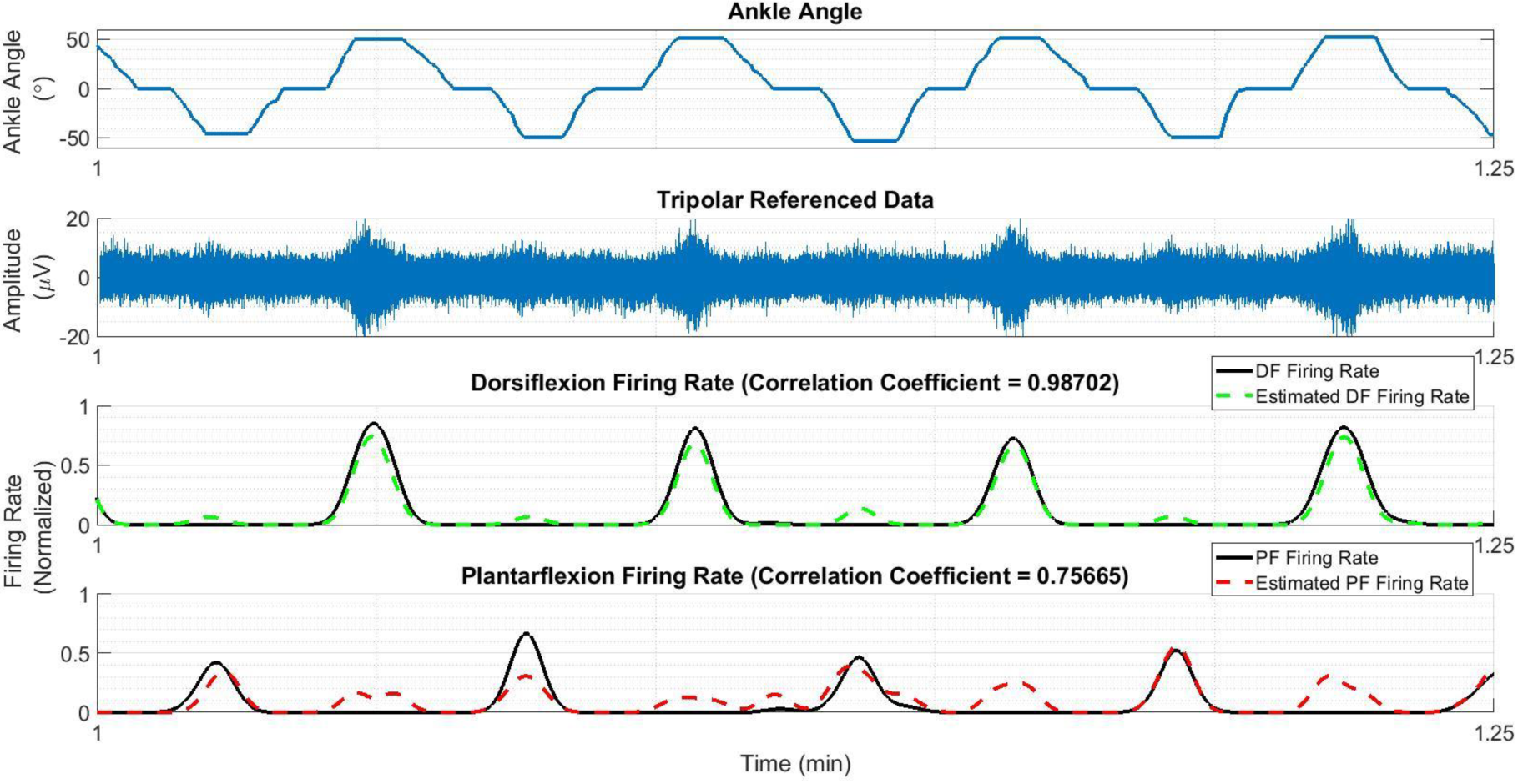
Relationship of ankle angle to the mean tripolar signal of the center ring and the estimated firing rates, obtained using the neural net classifier followed by removal of spurious CAPs using an optimal threshold. Plots are created from the recordings from Rat 6.

## 4. Discussion

The classification accuracies and firing rate reconstructions achieved in this study support the feasibility of CAP-based classification from multi-contact nerve cuff electrodes. However, further work is needed to improve the selectivity before robust and reliable CAP-based classification is achievable. Key factors determining the performance limitations of this method are discussed below.

The difference in the conduction velocities of the pathways being monitored is likely to have an influence on classification performance. In our previous simulation study [5], we examined the selectivity achievable with spatiotemporal templates, and found that this method could outperform beamforming [33] and VSR methods [26-28] for CAP-based classification. The performance improvement was greatest when the neural pathways were located in different fascicles and had different conduction velocities. In the present study, the experimental protocol activated neural pathways located in different fascicles and having fibers of similar conduction velocities (70.06 ± 12.26, and 71.93 ± 16.96 m/sfor the tibial and peroneal fascicle for the dorsiflexion and plantarflexion proprioceptive stimuli respectively, and 56.79 ± 10.34 m/s for the sural fascicle for the touch stimuli).

It was therefore not possible to directly examine how significant differences in conduction velocities would impact the classification. On the other hand, the *in vivo* results are close to the previous simulation results for the case of similar fiber types in different fascicles (F_1_-scores of 0.446 ± 0.157 in the *in vivo* results here vs. 0.412 ± 0.021 in the closest matching case in [5], using an estimated SNR of -10 dB). This comparison validates our previous simulations. It is therefore reasonable to conjecture that the performance *in vivo* would increase when attempting to distinguish pathways with more distinct conduction velocities, as suggested by the simulation results in [5].

Multiple aspects of our results point to SNR as the major contributor to the classification performance. The classification results in Figure 4 compare the discrimination potential of the different CAP-based classifiers to the RF-RBI approach. The classifiers show adequate discriminability while the RF-RBI approach was able to provide high mean F_1_-scores as the bin size increases. These findings are consistent with studies that use RBI for preprocessing before classifying or analyzing different activities/stimulus [32-34, 43, 44]. The improvements in SNR obtained from the RBI process are clearly beneficial for discrimination, as expected. These improvements come at a cost of decreased temporal resolution and potentially greater challenges in discriminating pathways with activity that overlap in time.

The impact of SNR is shown more directly through Figure 5. As the SNR increases, the mean F_1_-scores also increases which is most notably seen with the neural net classifier. The mean F_1_-scores increased from 0.539 ± 0.138 at SNR observed *in vivo* to 0.725 ± 0.150from an increase in SNR by a factor of 2 (average of 4 CAPs). Recall that the F_1_-score reported in Figure 4 for the SNR observed *in vivo* was computed using classifiers trained without data augmentation, in order to solely quantify the effects of SNR on F_1_-score. Hence, the performance shown for this SNR in Figure 5 is slightly lower than that shown in Figure 4, where data augmentation was used. The results shown in Figure 5 suggest that CAP-based classification from multi-contact nerve electrodes will be viable if low-noise recordings can be obtained, for example using hardware averaging [45].

In this study, certain factors were not optimized that may have contributed to a lower SNR. SNR could have been improved by adjusting the electrode surface contact with the nerve (the same electrode dimensions were used for all rats), by using different surgical techniques (e.g. by suturing closed the incision point), and by minimizing noise from cables and instrumentation [34] as well as shielding of the data acquisition equipment. All of these different factors could contribute to improved classification leading to greater discriminability of neural pathways.

Ultimately, the accurate classification of individual CAPs is of interest insofar as it can be used to estimate the firing patterns of the target pathways. Therefore, occasional CAP-based misclassifications are not worrisome if they do not have a large impact on the firing rate estimates.

The Pearson correlation coefficients found when the threshold (Eq. 1) was used showed robust tracking of one of the 2 activities (dorsiflexion or plantarflexion). The other activity’s firing rate was not as reliably tracked. The Pearson correlation coefficients increased when spurious CAPs were removed using a threshold on the outputted probabilities and allowed for both activities to be tracked more robustly. This suggests that reliable detection of CAPs before classification also plays an important role in the ability to track firing rates.

Thus, the firing rate reconstruction appears to be determined by two factors. Firstly, the classification accuracy which determines how well each CAP can be assigned to the correct neural pathway. Secondly, the CAP detection which reflects how well CAPs can be extracted from the noise. If CAPs from a given pathway cannot be reliably detected, the reconstructed firing rate for that pathway will be biased. Based on these observations, the use of a reliable CAP detection algorithm will be crucial to the viability of a CAP-based classification approach. It is important to note that the use of a delay-and-add or related operation can be of significant help in this regard, such that CAPs do not need to be detected directly in the noisy raw nerve cuff recordings.

The relationship illustrated in Figure 7 provides additional evidence that the reconstructed firing rates are related to the proprioceptive activity, and thus are encoding functional information. While inaccuracies remain (particularly for plantarflexion in this example), these are in line with the classification performance found in the rest of the analysis. Joint angle prediction from the firing rates is of interest in closed-loop functional neuromuscular stimulation systems [46-48] and other neuroprosthetic applications, though outside the scope of this study.

Both machine learning approaches were found to outperform the matched filter approach on the individual CAP classification tasks. The matched filter approach uses a template obtaining by averaging multiple CAPs, and correlates this template with an unknown signal. The template can be affected by spurious CAPs resulting in lower classification accuracy. In addition, the templates for neural pathways of similar fiber types may have only subtle differences, difficult to detect by this algorithm. The machine learning approaches on the other hand were trained directly on individual CAP examples, and are able to emphasize discriminating features more effectively than the matched filter. The RF-RBI algorithm provided the best classification results likely due to the RBI windowing technique, which greatly enhances the SNR. However, this comes at the cost of temporal resolution, which is an advantage of the CAP-based approach.

While the present work demonstrates the possibility of CAP-based classification, several factors may have affected the results in this *in vivo* study. Factors such as the positioning of the cuff electrode, the size of the nerve, or the amount of electrode surface contact on the nerve (nerve too small or large for the electrode) must be studied more closely to quantify their effects on the classification results. The manual delivery of the stimulus may have also introduced variability in the data set. Furthermore, being an acute study, electromyogram (EMG) interference from muscles are not seen in the recordings, and long-term changes in the recordings over time are not reflected. A chronic study must be performed where these effects can be studied in order to further assess the feasibility of this CAP-based classification approach.

The CAP labels used as the ground truth were obtained from the synchronized video data showing the stimulus applications, rather than from direct neural recordings in the tibial, peroneal and sural fascicles. Although much care was taken to selectively activate each fascicle (e.g. dorsiflexion using clip on the claws to avoid cutaneous input), there is a possibility that there was contamination within each activity. Labeling of dorsiflexion and plantarflexion activity may have also contained some mislabeled CAPs, which would introduce some inaccuracies. CAPs were labelled as dorsiflexion or plantarflexion based on the position of the ankle with respect to the neutral angle in the video recording, rather than based on the direction of movement. However, as seen in [38], neural activity from the return to neutral position from full dorsi- or plantar- flexion produces little neural activity in the unintended pathway. In addition, to verify that labelling was appropriate, the performance for 2 rats in which CAPs were labeled by direction of movement rather than position were compared to the results reported above and the classification accuracy did not increase (2-class CAP-based classification with the neural network, results not shown). Lastly, the high performance of the RF-RBI method demonstrates that good class separability exists within our dataset. Therefore, while the metrics reported in this study may be slightly affected by the factors describe above, we do not expect that the overall trends observed would change.

In practice, the need for calibration would be a disadvantage requiring acquisition of the CAP spatiotemporal signatures of pathways of interest and training of the classifiers. This tradeoff is not unreasonable if CAP-based classification is successful, which would result in the ability to reconstruct firing rates at finer time resolutions and improvement in selective recording performance in situations where multiple neural pathways are active over a given period of time.

This study suggests that classification of *individual, naturally evoked* CAPS using multi-contact nerve cuffs is possible. Further refinements to the classification algorithms, combined with improvements to the SNR and CAP detection, could likely yield robust discrimination performance and improved tracking of neuronal firing rates. These firing rates could be used to predict joint angles and other parameters of interest providing a significant step towards realizing applications such as an intuitive bi-directional control of assistive devices (i.e. prosthetic limb) that are able to produce more natural movements.

## Acknowledgements

This work was supported in part by the Natural Sciences and Engineering Research Council of Canada (RGPIN-2014-05498, RGPIN-2016-06329 and the NSERC Alexander Graham Bell Canada Graduate Scholarship-Doctoral Program) and the Institute of Biomaterials and Biomedical Engineering at the University of Toronto. The authors would also like to thank Jessica Trac for her assistance in the collection of the *in vivo* data.

## Author Contributions

All authors developed the algorithms. R.K and J.Z conceived the experiment. R.K and J.Z conducted the experiments. R.K. analyzed the results. R.K. and J.Z wrote the manuscript. All authors reviewed the manuscript.

